# The structure of the neurotoxin palytoxin determined by MicroED

**DOI:** 10.1101/2023.03.31.535166

**Authors:** Cody Gillman, Khushboo Patel, Johan Unge, Tamir Gonen

## Abstract

Palytoxin (PTX) is a potent neurotoxin found in marine animals that can cause serious symptoms such as muscle contractions, haemolysis of red blood cells and potassium leakage. Despite years of research, very little is known about the mechanism of PTX. However, recent advances in the field of cryoEM, specifically the use of microcrystal electron diffraction (MicroED), have allowed us to determine the structure of PTX. It was discovered that PTX folds into a hairpin motif and is able to bind to the extracellular gate of Na,K-ATPase, which is responsible for maintaining the electrochemical gradient across the plasma membrane. These findings, along with molecular docking simulations, have provided important insights into the mechanism of PTX and can potentially aid in the development of molecular agents for treating cases of PTX exposure.

## Introduction

Na,K-ATPase is an essential protein for maintaining proper cell function and is targeted by the potent marine toxin palytoxin (PTX) (Habermann, 1989; Christian Skou & Esmann, 1992; Tubaro *et al*., 2012). PTX is a non-proteinaceous natural product that was first isolated from tropical marine corals and later found in dinoflagellates(Usami *et al*., 1995; Taniyama *et al*., 2003; Ukena *et al*., 2014; Moore & Scheuer, 1971). PTX can accumulate to dangerous levels in seafood, leading to serious illness and death in those who consume it (Patocka *et al*., 2015; Deeds & Schwartz, 2010; Fukui *et al*., 1987; Rhodes *et al*., 2002). Aquarium hobbyists may also be exposed to PTX when mishandling Palythoa coral or inhaling aerosolized PTX (Hoffmann *et al*., 2008; Rumore & Houst, 2014). Inhaling PTX from blooming events of *Ostreopsis* has also caused severe illness and hospitalization (Ciminiello *et al*., 2006). Understanding the structure of PTX and how it binds to Na,K-ATPase is crucial for developing molecular agents that can treat cases of PTX exposure and protect against its toxic effects.

PTX ‘s binding to Na,K-ATPase with high affinity and its ability to convert it into a passive cation pore has serious implications for cellular function and can lead to a range of health effects, including skeletal muscle contractions, heart failure, hemolysis, and platelet aggregation (Böttinger *et al*., 1986; Riobó & Franco, 2011; Artigas & Gadsby, 2003; Wang & Horisberger, 1997). The irreversible depolarization of the membrane caused by PTX can also contribute to bone resorption and tumorigenesis (Lazzaro *et al*., 1987; Aligizaki *et al*., 2011). The extremely low lethal dose for humans highlights the severity of PTX poisoning (Tubaro *et al*., 2011; Wiles *et al*., 1974).

The development of anti-PTX molecules that can inhibit the binding of PTX on Na,K-ATPase is crucial for the treatment of PTX exposure. The structure of PTX when bound to an antibody fragment (scFv) was determined using microcrystal electron diffraction (MicroED) (Shi *et al*., 2013; Nannenga *et al*., 2014)at 3.2 Å resolution. This provided valuable information on the binding mode of PTX, which was then used to perform docking simulations to determine the potential binding mode of PTX on Na,K-ATPase. These findings pave the way for the development of molecular agents that can treat cases of PTX exposure by inhibiting the binding of PTX on Na,K-ATPase, and can potentially save many lives.

## Results

### Characterization of scFv-PTX complex and crystallization

To date, very little has been uncovered about the three-dimensional structure of PTX. Many studies have utilized anti-PTX antibodies to investigate PTX (Lau *et al*., 1995; Taniyama *et al*., 2003; Levine *et al*., 1987). The scFv antibody used in this study is a 26 kDa protein developed, expressed, and purified by Zabbio (San Diego, CA). The binding of PTX to scFv was confirmed using size exclusion chromatography (SEC). The shift in the SEC trace of free scFv and PTX-bound scFv was compared to confirm binding (Figure 1A). Furthermore, the binding affinity of PTX to scFv was determined using microscale thermophoresis (MST). PTX binds to scFv at a K_D_ of 2.1 μM (Figure 1B).

**Figure 1.**
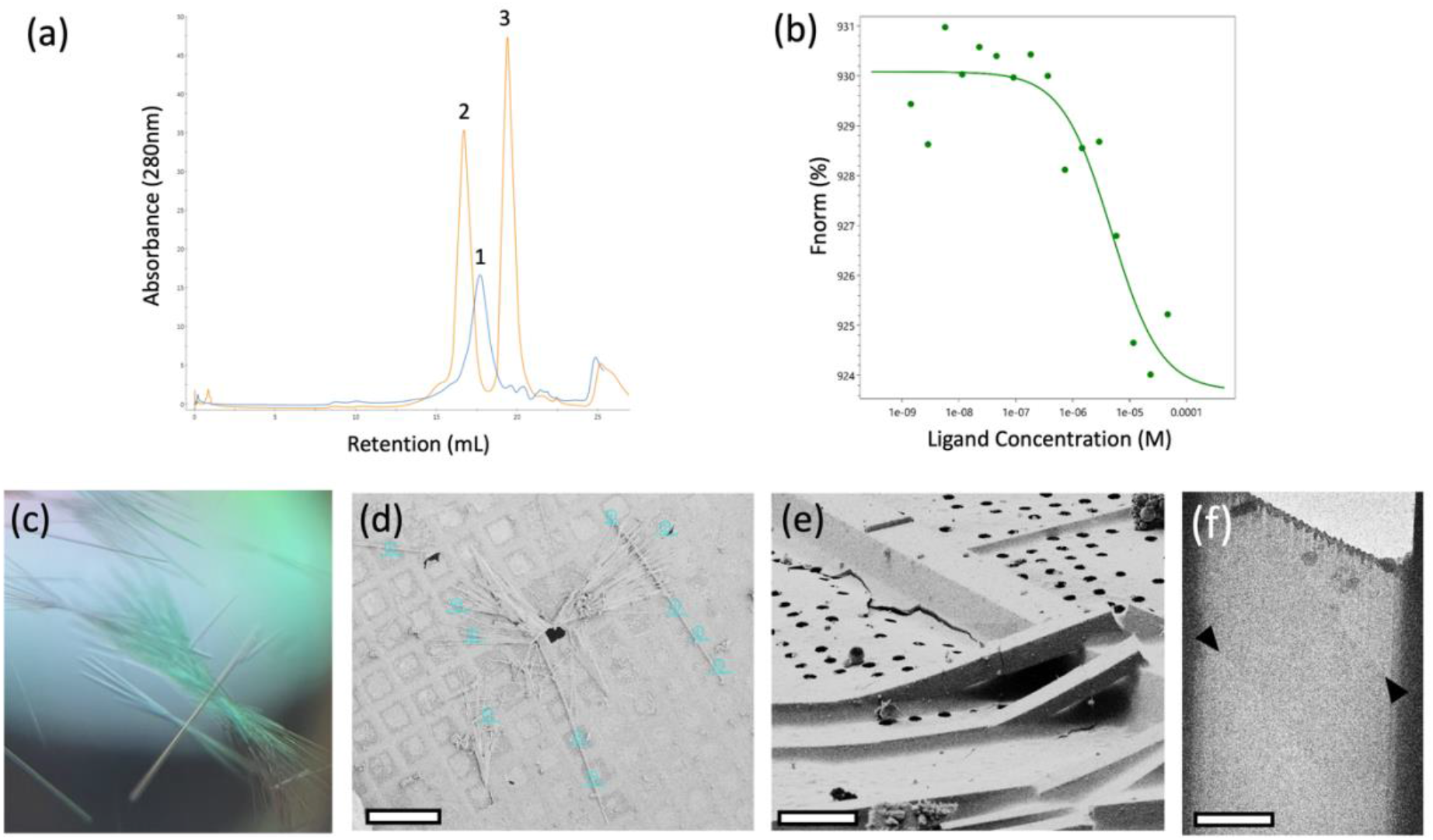
Crystallization of scFv-PTX. (a) Overlay of size-exclusion chromatography traces of scFv alone (blue trace) and scFv-PTX complex (orange trace). (b) Microscale thermophoresis binding assay confirming binding between scFv-PTX complex. This assay indicates that palytoxin binds to ScFv with K_D_ = 2.1 μM affinity (c) Light microscope image of scFv-PTX complex crystals (d) SEM image of crystals viewed normal to the grid support surface prior to FIB milling. Scale bar = 200 µm. (e) FIB image of a crystal milling site. Scale bar = 10 µm. (f) Lamella imaged normal to the grid surface in the TEM after milling. Scale bar = 10 µm. The lamella was 200nm thick.

SEC fractions corresponding to the stable scFv-PTX complex were collected, concentrated to 10 mg/mL, and subjected to sparse matrix crystallization screening to identify crystallization condition hits. The scFv-PTX complex was crystallized by the hanging drop vapor diffusion technique. The well solution contained 27 % Jeffamine ED-2001 pH 7.0 and 100 mM sodium citrate tribasic dihydrate pH 5.6. The scFv-PTX complex was combined with the well solution at 2:1 (v/v) ratio. The crystals were thin rods that formed in dense bundles (Figure 1C). The average size of each crystal was 5 μm x 500 μm. The crystals in the drop were then transferred to an electron microscopy (EM) sample grid, blotted to remove surrounding crystallization media, and vitrified by plunge freezing into liquid ethane. Crystals were stored in liquid nitrogen prior to use.

### Preparaing crystal lamellae and collection of MicroED data

The crystals that were obtained for this study were too thick for MicroED and needed to be thinned to a thickness that would allow for the transmission of electrons (Martynowycz *et al*., 2021). To achieve this, thin crystal lamellae were produced using a cryogenic focused ion beam scanning electron microscope (FIB/SEM) milling instrument (Martynowycz *et al*., 2019). The process began by loading the EM grid with crystals into the FIB/SEM at cryogenic temperature, followed by imaging using the SEM (Figure 1D). Potential milling sites were then observed in the FIB view of the specimen (Figure 1E), and the targeted crystal and surrounding media were milled into a thin lamella using the gallium ion beam. The final product was a lamella that measured 7 μm wide and 300 nm thick.

After the crystal lamellae were produced, they were transferred to a Titan Krios transmission electron microscope that was operating at 300 kV and cooled cryogenically. The sites of the lamellae were identified using low magnification imaging and adjusted to eucentric height. To ensure high-resolution diffraction, a preview of the lamella was taken (Figure 1F and 2A). The data was collected using continuous rotation MicroED (Nannenga *et al*., 2014), with a Falcon4 direct electron detector set to counting mode. The highest resolution spots were observed at 3.2 A resolution.

### Determining the MicroED structure of scFv-PTX complex

MicroED data were converted to standard crystallographic formats using software freely available on our website (https://cryoem.ucla.edu/microed). The data were indexed and integrated in XDS (Kabsch, 2010*b*) to a resolution of 3.2 Å, which corresponds to where the CC1/2 was approximately 32%. Reflection data from three crystal lamellae were merged to increase completeness. Phases for the MicroED reflections were determined by molecular replacement using scFv 4B08 (PDB 5YD3) (Miyanabe *et al*., 2018) as a search model. scFv 4B08 and scFv have 45% sequence identity. The space group was determined to be P4_1_2_1_2 and unit cell dimensions were 69.95, 69.95, 289.48 (a, b, c) (Å) and 90, 90, 90 (α, β, γ) (°). The structure was refined using electron scattering factors (Table 1) using phenix.refine (Afonine *et al*., 2012). The scFv-PTX complex is in dimeric form with one PTX bound to each scFv monomer. The density map contoured at 1.5 σ had continuous density for the backbone of the scFv and the side chains of the amino acids were also well resolved (Figure 2B). Continuous density was obtained for PTX after multiple rounds of refinement (Figure 2C). The R_work_ and R_free_ of the refinement were 28% and 32%, respectively.

**Table 1.** MicroED structure statistics of scFv-PTX complex.

| Data | Parameter | Measure |
| --- | --- | --- |
| Data Collection | Accelerating voltage (kV) | 300 |
|  | Electron source | field emission gun |
|  | Total accumulated exposure (e-/Å<sup>2</sup>) | 0.64 |
|  | Microscope | Thermo Fisher Titan Krios |
|  | Camera | Falcon 4 electron counting |
|  | Rotation rate (deg/sec) | 0.2 |
|  | Wavelength (Å) | 0.019687 |
| Data analysis | No. of crystals | 3 |
|  | Resolution range (Å) | 46.80-3.20 |
|  | Space group | P4<sub>1</sub>2<sub>1</sub>2 |
|  | a, c, c (Å) | 69.95, 69.95, 289.48 |
|  | α, β, γ (°) | 90, 90, 90 |
|  | Reflections, total/unique | 260311/12734 |
|  | Multiplicity | 20.44 |
|  | Completeness (%) | 99.3 |
|  | Mean I/σ(I) | 2.59 |
|  | CC1/2 | 95.3 |
|  | Rwork | 0.2830 |
|  | Rfree | 0.3229 |

**Figure 2.**
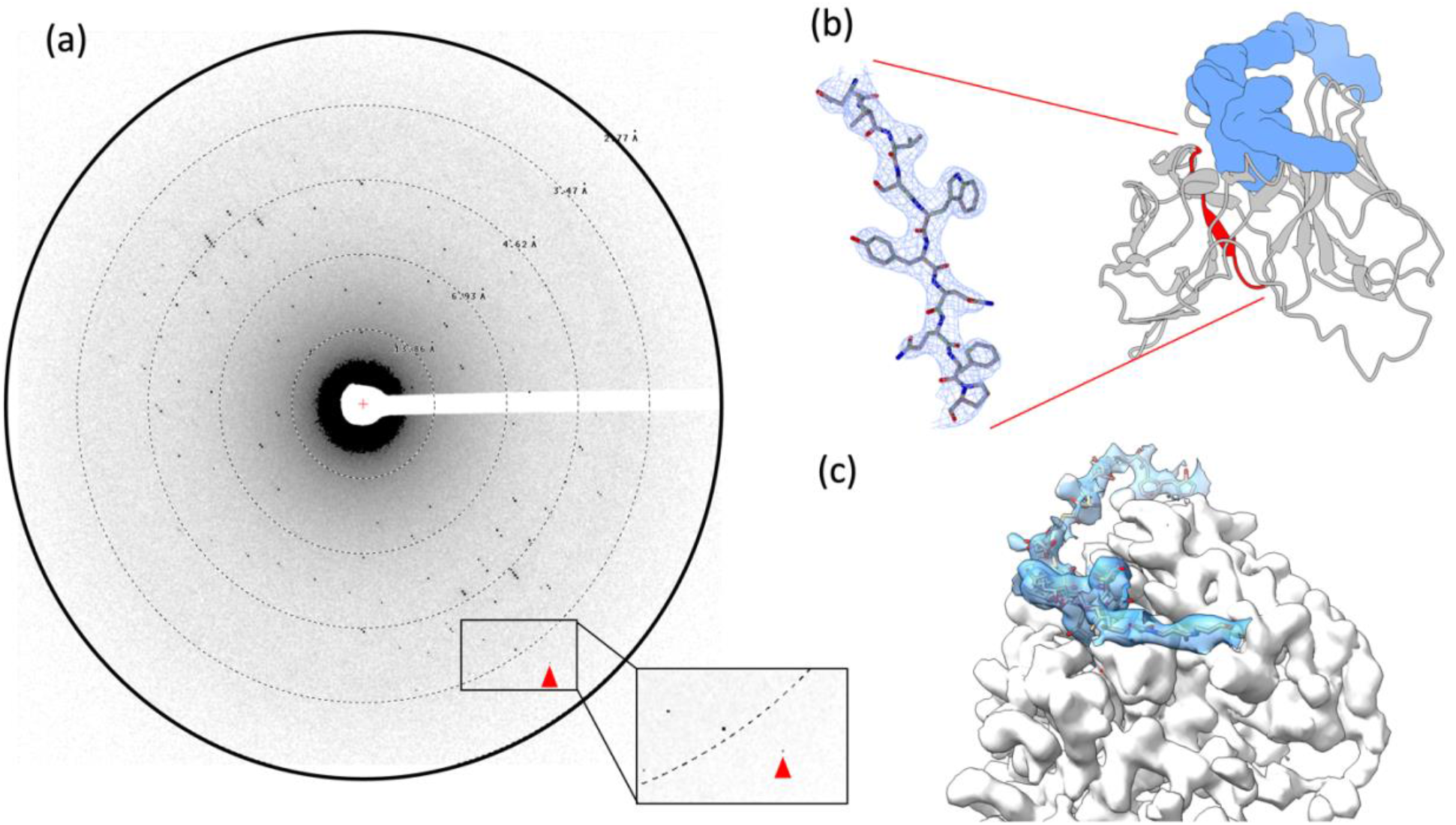
The MicroED structure of PTX in complex with scFv. (a) Representative MicroED image. The highest resolution spots were observed at 3.2 Å (red arrowhead). (b) Structure of scFv-PTX complex. The scFv is represented as a cartoon in grey and PTX is represented as a surface model in blue. A beta-strand was selected (red) to provide a sample of the 2mFo–DFc map (blue mesh), which was contoured at 1.5 σ with a 2-Å carve. (c) The overall 2mFo–DFc map of density for both PTX (solid blue) and scFv (solid white) in complex. The density map was contoured at 1.5 σ.

### The MicroED structure of scFv-PTX complex

The scFv creates a binding pocket into which an internal segment of the PTX chain is inserted, forming a hairpin motif (depicted in Figure 3). The deepest part of the pocket consists of a double ring with two cyclic ethers, which is flanked by two hydrocarbon chains running antiparallel to each other. Hydrogen bonds are formed between a cyclic ether of the double ring and residues Y106, E108, and Y169 of the scFv. Hydrophobic amino acid side chains, including F59, W224, V231, and V165, sequester the hydrophobic segments of PTX flanking the double ring from the external aqueous environment. At the entrance to the binding pocket, a network of intramolecular hydrogen bonds is formed by several hydroxyl groups. Outside the binding pocket, the two tail ends of PTX are observed traveling in opposite directions in the solvent channel of the crystal.

**Figure 3.**
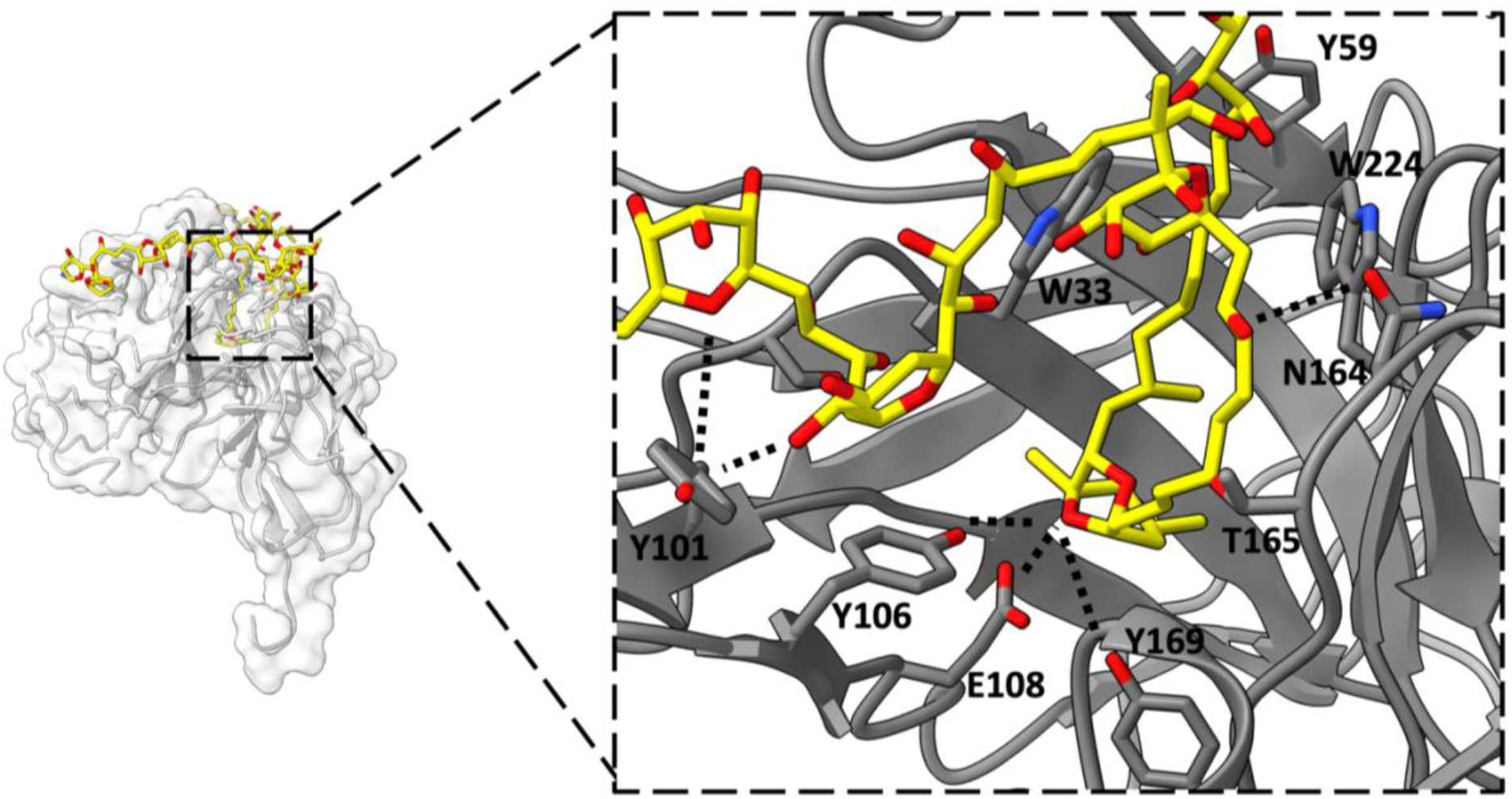
Binding interactions between scFv and PTX. An internal region of PTX (represented as yellow carbons and heteroatoms) folds into a hairpin motif and inserts into the scFv binding pocket. A double ring of PTX forms several H-bonding interactions with polar amino acid side chains at the deepest part of the scFv binding pocket (Y101, Y106, E108). A belt of hydrophobic amino acid side chains interact with the hydrophobic chains of PTX (V165, V231, W224, F59). Additional H-bond interactions take place near the mouth of the binding pocket (N164, W33, Y101).

**Figure 4:**
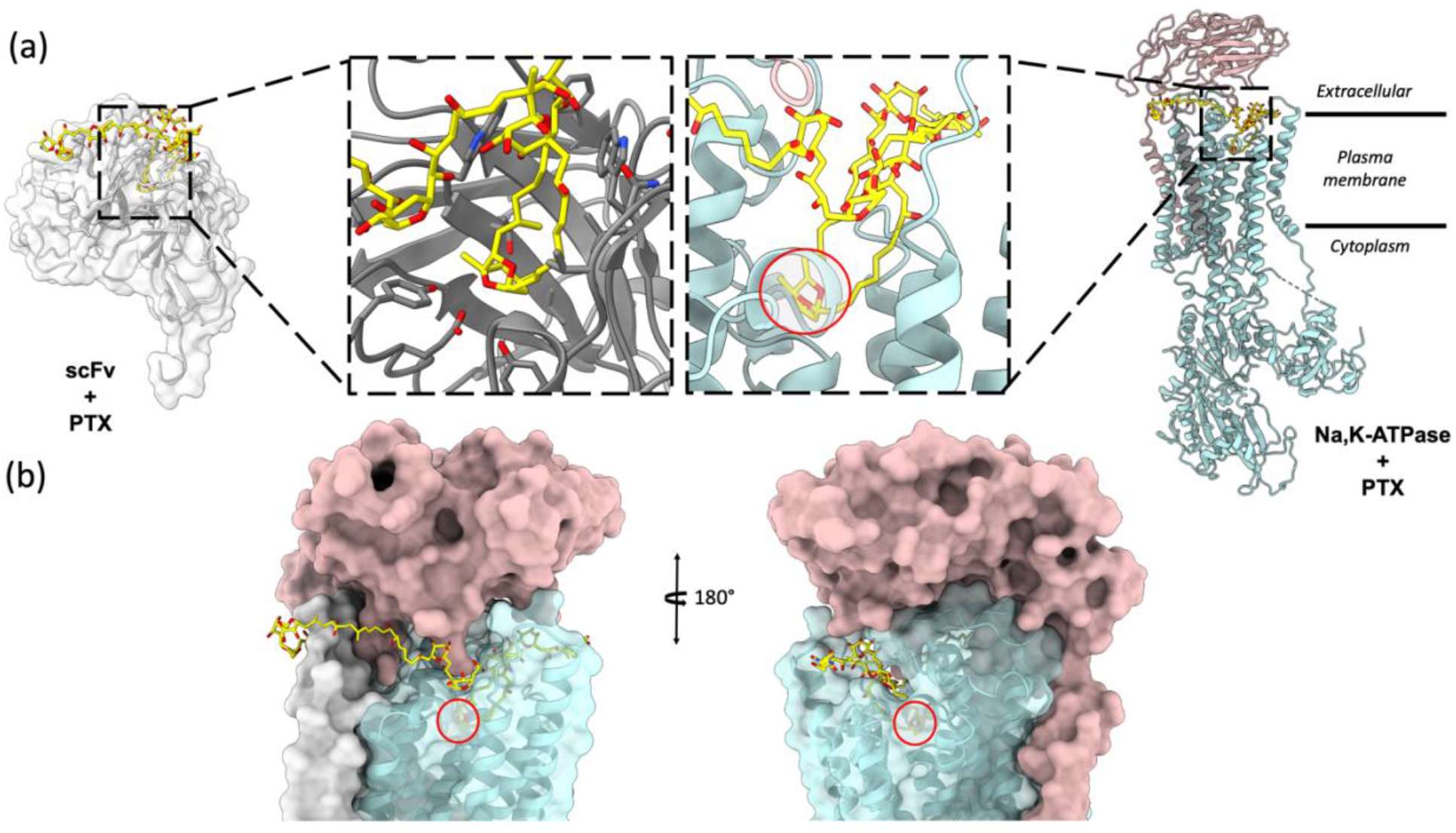
Molecular docking of PTX to Na,K – ATPase. (A) Comparison of binding modes of PTX to scFv and Na,K – ATPase. The hairpin motif of PTX shown as yellow sticks interacts with both scFv shown as white surface and Na, K – ATPase shown as cyan and salmon pink ribbons. (B) Surface representation of Na,K-ATPase shows that the hairpin motif of PTX binds the extracellular gate of the pump in a plug-like manner.

**Figure 5:**
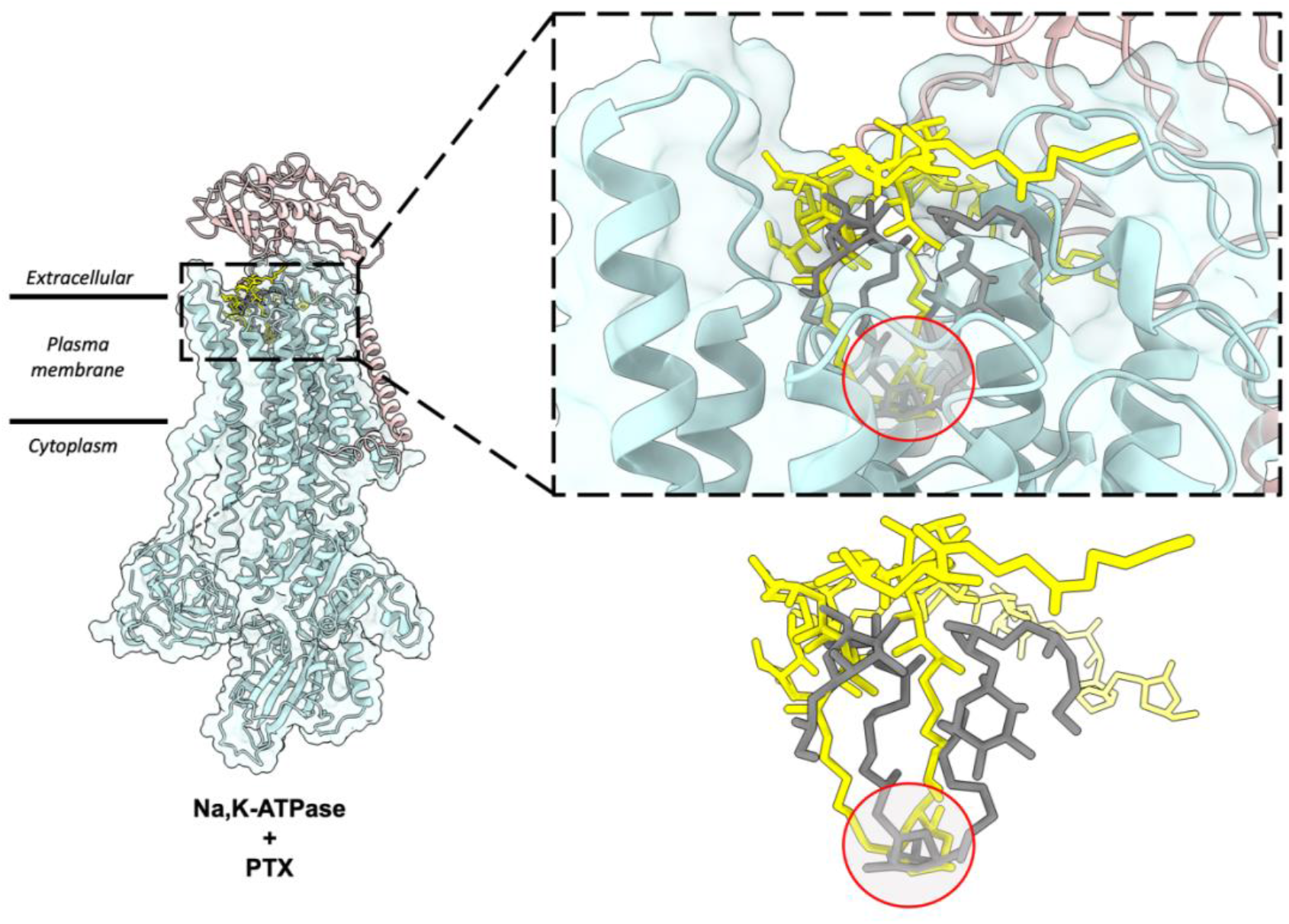
Comparison of top docking solutions from patchdock and autodock vina. The patchdock solution (yellow sticks) and vina solution (grey sticks) show that the finger motif of PTX binds to the extracellular gate of the Na,K-ATPase. In both solutions, the double ring (highlighted by red circle) interacts with Na,K-ATPase by forming hydrogen bonds and the surrounding helices form hydrophobic interactions with the hydrocarbon chains of the hairpin motif.

### Molecular docking

The scFv-PTX complex structure obtained by MicroED was used to investigate the potential binding of PTX to the Na,K-ATPase protein using molecular docking. The hairpin-like motif of PTX from the scFv-PTX complex was used as the ligand, and the human Na,K-ATPase structure in E1 state was used as the receptor protein molecule (Guo *et al*., 2022). Rigid docking simulations were performed using the Patchdock server (Schneidman-Duhovny *et al*., 2005), which suggested that the hairpin motif of PTX binds to the extracellular gate of the Na,K-ATPase protein. The cyclic ether forms hydrogen bonds, while the hydrocarbon chains are protected by hydrophobic transmembrane alpha-helices. The two tails of PTX are between the alpha and beta subunits of Na,K-ATPase. Flexible docking simulations were performed using Autodock Vina (Trott & Olson, 2009) to further confirm the Patchdock results. The results indicated that the hairpin motif of PTX could bind to the similar binding pocket in the Na,K-ATPase, forming a plug-like structure that blocks the channel, thereby rendering it inactive. This is consistent with previously reported functional assays (Vale & Ares, 2007; Ramos & Vasconcelos, 2010). Both binding simulations suggest that PTX ‘s hairpin motif could potentially bind to the Na,K-ATPase and block the channel, leading to a possible explanation for the cytotoxic effects of PTX.

## Discussion

The findings of this study provide significant insights into the molecular mechanism of PTX binding to Na,K-ATPase. The hairpin motif formed by the hydrophobic region of PTX when bound to scFv was found to also fit into the extracellular gate of Na,K-ATPase like a plug, blocking the ion channel and rendering the pump inactive. This corroborates earlier reports that this region of PTX is chiefly responsible for its interaction with biomembranes and may be important in the conversion of Na,K-ATPase from a pump to a passive cation pore (Harmel & Apell, 2006; Habermann *et al*., 1981). The molecular docking simulations performed in this study provide a model of how PTX binding to Na,K-ATPase takes place and the results suggest that PTX likely adopts the same hairpin fold when bound to Na,K-ATPase. Overall, this study represents an important step towards a better understanding of the molecular mechanisms involved in PTX binding and its effects on Na,K-ATPase.

The use of scFv in this study allowed for the determination of the 3D structure of the scFv-PTX complex using MicroED. The scFv-PTX complex was crystallized using the hanging drop vapor diffusion method, and the crystals were thinned using cryogenic FIB milling prior to MicroED diffraction. The needle-shaped crystals of the scFv-PTX complex that were obtained for this study were typically thin (3-5 µm) and long (several hundred microns) and they formed in bundles. Such morphologies are extremely challenging for analyses by x-ray crystallography, often leading to multiple lattices and weak scattering. Using MicroED and FIB milling was advantageous in this case because the entire crystal bundle could be transferred to the EM grid and crystal sites were accessed by using a FIB mill to generate crystal lamellae and ultimately a MicroED structure.

The structure showed that PTX binds to the scFv in a highly specific manner, forming several key interactions with amino acid residues in the complementarity-determining regions (CDRs) of the scFv. In particular, PTX binds to the CDR H3 loop of the scFv, which is known to be a critical region for antigen binding (Figure 2B). The 3D binding mode of PTX to scFv was used to perform docking simulations to predict the binding mode of PTX to Na,K-ATPase. The docking simulations suggested that PTX binds to Na,K-ATPase in a similar manner to scFv, with the key interactions occurring in the extracellular ion gate of Na,K-ATPase (Figure 3). This provides important insights into the mechanism of PTX binding to Na,K-ATPase and can aid in the development of anti-PTX molecules that prevent the binding of PTX to Na,K-ATPase.

This research paves the way for the development of possible treatments for PTX exposure. The detailed structural information obtained from our MicroED study can aid in the creation of new inhibitors that can block the binding of PTX to Na,K-ATPase, thus preventing its toxic effects. Additionally, the knowledge gained from this study can be applied to develop methods for identifying and monitoring the accumulation of PTX and its analogues in the environment, potentially preventing harmful exposure to both humans and marine life. In summary, this research not only elucidates the mechanism of action of PTX, but also offers valuable insights into the development of potential therapeutics and environmental monitoring techniques. Significantly, this study reinforces the utility of MicroED as a powerful tool for revealing the structures of important biomolecules, such as the long-awaited structure of palytoxin, which has been elusive to x-ray crystallography.

## Acknowledgments

The authors would like to thank Zabbio (San Diego, CA) for development and generation of ScFv. This study was supported by the National Institutes of Health P41GM136508 and the Department of Defense HDTRA1-21-1-0004. The Gonen laboratory is supported by funds from the Howard Hughes Medical Institute. Coordinates and maps were deposited in the protein data bank (Accession code XXXX) and the EM Data bank (Accession code YYYY).

## Methods

### Materials

All reagents were made with MilliQ water. Palytoxin was purchased from Fuji Film (Japan). Crystallization reagents were purchased from Hampton Research (Aliso Viejo, CA). Monolith protein labeling kit RED-NHS 2nd generation was purchased from NanoTemper Technologies (Munich, Germany).

### Microscale thermophoresis

The anti-PTX scFv was prepared in 25 mM HEPES (pH 7.4), 150 mM NaCl at a concentration of 13 μM, and labeled with RED-NHS dye (NanoTemper). PTX was prepared in 25 mM HEPES (pH 7.4), 150 mM NaCl at a concentration of 93 μM. Labeled scFv was diluted to 80 nM and added in a 1:1 ratio to a dilution series of 46.5 μM down to 4.65 × 10^-15 μM of PTX and 0 μM of PTX. Mixtures were loaded into premium capillaries (Monolith Capillaries, NanoTemper Technologies). Thermophoresis was measured at 21°C for 15 sec with 50% LED power and 100% (auto-detect) power.

### Crystallization

The complex was purified by size-exclusion chromatography and the elution fractions were concentrated to 10 mg/mL. Palytoxin was incubated with scFv at a 2:1 molar ratio for 30 minutes at room temperature. A sparse matrix screening hit was identified in the PEGRxHT well condition C01 (Hampton Research) by sitting drop vapor diffusion using a Mosquito crystallization robot. This condition was optimized for robust crystallization using hanging drop vapor diffusion. In the final condition, the complex was crystallized by mixing with 27 % Jeffamine ED-2001 pH 7.0, and 100 mM sodium citrate tribasic dihydrate pH 5.6 in 1.5 uL drops with a 2:1 sample-to-mother liquor ratio.

### Cryo-preservation

The cover slip with crystal drop was removed from the screening tray and the drop was gently applied to a Cu200 R2/2 holey carbon EM grid (quantifoil). The EM grid was negatively glow-discharged prior to sample application. The grid was blotted in a Leica GP2 set to 95% humidity and 12°C and plunge-frozen into liquid ethane. The sample was stored in liquid nitrogen until further use.

### Machining crystal lamellae using the cryo-FIB/SEM

The vitrified EM grid was loaded into a Thermo Fisher Aquilos dual-beam FIB/SEM operating at cryogenic temperature following established procedures (Martynowycz *et al*., 2019). The sample was sputter coated with a thin layer of platinum to preserve the sample during imaging and ion beam milling. A whole-grid atlas of the drop was acquired by the SEM and potential milling sites were selected. The targeted crystal and surrounding media were milled into a thin lamella using the gallium ion beam. The first stage of milling used a beam current of 0.5 nA and gradually decreased to a minimum of 10 pA as the lamella became thinner at later stages of milling. The final lamellae were 7 μm wide and 200 nm thick.

### MicroED Data Collection

Grids with milled lamellae were transferred to a cryogenically cooled Thermo Fisher Scientific Titan Krios G3i TEM operating at an accelerating voltage of 300 kV. The Krios was equipped with a field emission gun and a Falcon4 direct electron detector. A low magnification atlas of the grid was acquired using EPUD (Thermo Fisher) to locate milled lamellae. The stage was translated to the lamellae position and the eucentric height was set. The 100 μm selected area aperture was inserted and centered on the crystal to block background reflections. In diffraction mode, the beam was defined using a 50 μm C2 aperture, a spotsize of 11, and a beam diameter of 20 μm. MicroED data were collected by continuously rotating the stage at 0.2 ° / s. MicroED data from three different crystal lamellae were selected for downstream data processing.

### MicroED Data Processing

Diffraction movies in MRC format were converted to SMV format using MicroED tools (https://cryoem.ucla.edu/microed) (Martynowycz *et al*., 2019; Hattne *et al*., 2015). The diffraction dataset was indexed and integrated in *XDS* (Kabsh, 2010). Integrated intensities from three different crystal lamella were merged and scaled in *XSCALE* (Kabsch, 2010*a*).

### Structure solution and refinement

Phases for the MicroED reflections were determined by molecular replacement in PHASER (McCoy *et al*., 2007) using anti-Mcl1 scFv (PDB 6QF9) as the search model (Luptak *et al*., 2019). The solution was space group P4_1_2_1_2 and unit cell dimensions 69.95, 69.95, 289.48 (a, b, c) (Å) and 90, 90, 90 (α, β, γ) (°). The first refinement was performed with Coot and phenix.refine (Afonine *et al*., 2012) using isotropic B-factors and electron scattering factors. Occupancies were refined for alternative side chain conformations and SO_4_ and waters were manually placed during refinement. The final refinement used anisotropic B-factors, automatic water picking, and electron scattering factors and resulted in R_work_/R_free_ = 0.2830/0.3229 and resolution of 3.2 Å.

### Molecular docking

A human Na,K-ATPase structure in E1 state (PDB ID: 7E21) (Guo *et al*., 2022) without cofactors, waters and ligands was used as a receptor molecule and the 3D model of PTX from the scFv-PTX complex was used as a rigid ligand. For patchdock simulations, the PDB files of receptor and ligand molecules were submitted to the patchdock server (http://bioinfo3d.cs.tau.ac.il/PatchDock/php.php). The clustering RMSD was selected to be 4.0 and the complex type was selected to be protein-ligand complex. The results were emailed within 24h with a list of potential binding solutions of PTX numbered on the basis of geometric shape complementarity score. Higher complementarity scores indicate less possibility of steric clashes in the solution.

Full length PTX did not provide any solution when simulation was performed using Autodock vina, hence, a flexible fragment of PTX molecule consisting of only the hairpin motif was used to perform the binding simulations. The receptor and ligand were prepared using the MGL tools suite (https://ccsb.scripps.edu/mgltools/) and saved as pdbqt files. The receptor file contained partial charges and polar hydrogens. Any cofactors, waters and ligands were removed. For the ligand file, polar hydrogens were added and all the original torsion angles were kept intact. The receptor was treated as rigid but the ligand fragement was flexible. The simulations were run and the solutions were scored on the basis of binding energy (kcal/mol).

## Notes

### Competing Interest Statement

The authors have declared no competing interest.

### Summary of Updates

Updated methods section

## References

Afonine, P. V., Grosse-Kunstleve, R. W., Echols, N., Headd, J. J., Moriarty, N. W., Mustyakimov, M., Terwilliger, T. C., Urzhumtsev, A., Zwart, P. H. & Adams, P. D. (2012). Acta Crystallogr D Biol Crystallogr 68, 352–367.

Aligizaki, K., Katikou, P., Milandri, A. & Diogène, J. (2011). Toxicon 57, 390–399.

Artigas, P. & Gadsby, D. C. (2003). Proc Natl Acad Sci U S A 100, 501–505.

Böttinger, H., Béress, L. & Habermann, E. (1986). BBA - Biomembranes 861, 165–176.

Christian Skou, J. I. & Esmann, M. I. (1992). J Bioenerg Biomembr 24.

Ciminiello, P., Dell‘Aversano, C., Fattorusso, E., Forino, M., Magno, G. S., Tartaglione, L., Grillo, C. & Melchiorre, N. (2006). Anal Chem 78, 6153–6159.

Deeds, J. R. & Schwartz, M. D. (2010). Toxicon 56, 150–162.

Fukui, M., Murata, M., Inoue, A., Gawel, M. & Yasumoto, T. (1987). Toxicon 25, 1121–1124.

Guo, Y., Zhang, Y., Yan, R., Huang, B., Ye, F., Wu, L., Chi, X., shi, Y. & Zhou, Q. (2022). Nat Commun 13, https://doi.org/10.1038/s41467-022-31602-y.

Habermann, E. (1989). Toxicon 27, 1171–1187.

Habermann, E., Ahnert-Hilger, G., Chhatwal, G. S. & Beress, L. (1981). Biochimica et Biophysica Acta (BBA) - Biomembranes 649, 481–486.

Harmel, N. & Apell, H.-J. (2006). J Gen Physiol 128, 103–118.

Hattne, J., Reyes, F. E., Nannenga, B. L., Shi, D., De La Cruz, M. J., Leslie, A. G. W. & Gonen, T. (2015). Acta Crystallogr A Found Adv 71, 353–360.

Hoffmann, K., Hermanns-Clausen, M., Buhl, C., Büchler, M. W., Schemmer, P., Mebs, D. & Kauferstein, S. (2008). Toxicon 51, 1535–1537.

Kabsch, W. (2010a). Acta Crystallogr D Biol Crystallogr 66, 133–144.

Kabsch, W. (2010b). Acta Crystallogr D Biol Crystallogr 66, 125–132.

Kabsh, W. (2010). Acta Crystallogr D Biol Crystallogr 66, 125–132.

Lau, C. O., Tan, C. H., Khoo, H. E., Yuen, R., Lewis, R. J., Corpuz, G. P. & Bignami, G. S. (1995). Toxicon 33, 1373–1377.

Lazzaro, M., Tashjian Jr., A., Fujiki, H. & Levinef, L. (1987). Endocrinology 120, 1338–1345.

Levine, L., Fujiki, H., Gjika, H. B. & Van Vunakis, H. (1987). Toxicon 25, 1273–1282.

Luptak, J., Bista, M., Fisher, D., Flavell, L., Gao, N., Wickson, K., Kazmirski, S. L., Howard, T., Rawlins, P. B. & Hargreaves, D. (2019). Acta Crystallographica Section D 75, 1003–1014.

Martynowycz, M. W., Clabbers, M. T. B., Unge, J., Hattne, J. & Gonen, T. (2021). Proc Natl Acad Sci U S A 118, 1–7.

Martynowycz, M. W., Zhao, W., Hattne, J., Jensen, G. J. & Gonen, T. (2019). Structure 27, 545–548.e2.

McCoy, A. J., Grosse-Kunstleve, R. W., Adams, P. D., Winn, M. D., Storoni, L. C. & Read, R. J. (2007). J Appl Crystallogr 40, 658–674.

Miyanabe, K., Akiba, H., Kuroda, D., Nakakido, M., Kusano-Arai, O., Iwanari, H., Hamakubo, T., Caaveiro, J. M. M. & Tsumoto, K. (2018). The Journal of Biochemistry 164, 65–76.

Moore, R. E. & Scheuer, P. J. (1971). Science (1979) 172, 495–498.

Nannenga, B. L., Shi, D., Leslie, A. G. W. & Gonen, T. (2014). Nat Methods 11, 927–930.

Patocka, J., Gupta, R. C., Wu, Q. & Kuca, K. (2015). Journal of Huazhong University of Science and Technology [Medical Sciences] 35, 773–780.

Ramos, V. & Vasconcelos, V. (2010). Mar Drugs 8, 2021–2037.

Rhodes, L., Towers, N., Briggs, L., Munday, R. & Adamson, J. (2002). N Z J Mar Freshwater Res 36, 631–636.

Riobó, P. & Franco, J. M. (2011). Toxicon 57, 368–375.

Rumore, M. M. & Houst, B. M. (2014). Int J Case Rep Images 5, 501–504.

Schneidman-Duhovny, D., Inbar, Y., Nussinov, R. & Wolfson, H. J. (2005). Nucleic Acids Res 33, W363–W367.

Shi, D., Nannenga, B. L., Iadanza, M. G. & Gonen, T. (2013). Elife 2, e01345.

Taniyama, S., Arakawa, O., Terada, M., Nishio, S., Takatani, T., Mahmud, Y. & Noguchi, T. (2003). Toxicon 42, 29–33.

Trott, O. & Olson, A. J. (2009). J Comput Chem NA-NA.

Tubaro, A., Durando, P., Del Favero, G., Ansaldi, F., Icardi, G., Deeds, J. R. & Sosa, S. (2011). Toxicon 57, 478–495.

Tubaro, A., Sosa, S. & Hungerford, J. (2012). Toxicology and diversity of marine toxins. Veterinary toxicology: basic and clinical principles Academic press.

Ukena, T., Satake, M., Usami, M., Oshima, Y., Naoki, H., Fujita, T., Kan, Y. & Yasumoto, T. (2014). Biosci Biotechnol Biochem 65, 2585–2588.

Usami, M., Satake, M., Ishida, S., Inoue, A., Kan, Y. & Yasumoto, T. (1995). J Am Chem Soc 117, 5389–5390.

Vale, C. & Ares, I. R. (2007). Phycotoxins: Chemistry and Biochemistry 95–118.

Wang, X. & Horisberger, J. D. (1997). FEBS Lett 409, 391–395.

Wiles, J. S., Vick, J. A. & Christensen, M. K. (1974). Toxicon 12, 427–433.

